# Avoiding test set bias with rank-based prediction

**DOI:** 10.1101/005983

**Authors:** Prasad Patil, Pierre-Olivier Bachant-Winner, Benjamin Haibe-Kains, Jeffrey T. Leek

## Abstract

**Background:** Prior to applying genomic predictors to clinical samples, the genomic data must be properly normalized. The most effective normalization methods depend on the data from multiple patients. From a biomedical perspective this implies that predictions for a single patient may change depending on which other patient samples they are normalized with. This test set bias will occur when any cross-sample normalization is used before clinical prediction.

**Methods:** We developed a new prediction modeling framework based on the relative ranks of features within a sample in order to prevent the need for cross-sample normalization, therefore effectively avoiding test set bias. We employed modeling with previously published Top-Scoring Pairs (TSPs) methodology to build the rank-based predictors. We further investigated the robustness of the rank-based models in case of heterogeneous datasets using diverse microarray technologies.

**Results:** We demonstrated that results from existing genetic signatures which rely on normalizing test data may be unreproducible when the patient population changes composition or size. Using pairwise comparisons of features, we produced a ten gene, platform-robust, and interpretable alternative to the PAM50 subtyping signature and evaluated the robustness of our signature across 6,297 patients samples from 28 curated breast cancer microarray datasets spanning 15 different platforms.

**Conclusion:** We propose a new approach to developing genomic signatures that avoids test set bias through the robustness of rank-based features. Our small, interpretable alternative to PAM50 produces comparable predictions and patient survival differentiation to the original signature. Additionally, we are able to ensure that the same patient will be classified the same way in every context.

## Introduction

One of the most common barriers to the development and translation of genomic signatures is cross-sample variation in technology, normalization, and laboratories [1]. Technology, batch, and sampling artifacts have been responsible for the failure of genomic signatures [2, 3], unreproducibility of genomic results [4], and retraction of papers reporting genomic signatures [5]. Even highly successful signatures such as Mammaprint [6] have required platform-specific retraining before they could be translated to clinical use [7].

An underappreciated source of bias in genomic signatures is test set bias [8]. Test set bias occurs when the predictions for any single patient depend on the data for other patients in the test set. For example, suppose that the gene expression data for a single patient is normalized by subtracting the mean expression across all patients in the test set. Then the normalized value for any specific gene for that patient depends on the values for all the patients they are normalized with. The result is that a patient may get two different predictions using the same data and the same prediction algorithm, depending on the other patients used to normalize the test set data (Figure 1).

**Figure 1:**
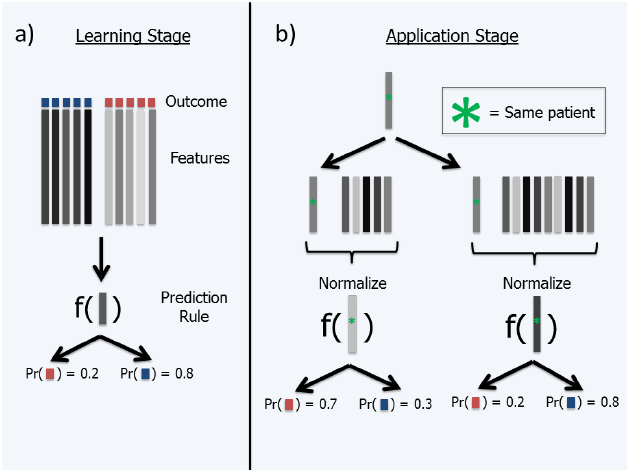
A description of how test set bias can alter class prediction for an individual patient. In panel a), we learn a model for predicting if a patient is in class R (red) or class B (blue). In our training data, the patients with darker grey features tend to be in class B, while the lighter patients are in class R. We develop a prediction rule from our training data and apply it to a new darker grey patient, we see that he is likely to be classified to class B. In panel b), we attempt to classify a single patient in the context of two different patient populations. We see that depending on the number and type of other patients in the population when we normalize the data, the resulting feature profile for our patient can be drastically different. This leads to different eventual classifications by our prediction rule. We contend that the ultimate classification of a patient should not depend on the characteristics of the test set, but rather solely on the characteristics of the patient himself.

Some normalization methods [9, 10] and some batch correction methods [11, 12] have addressed this issue by normalizing each sample against a fixed, or “frozen”, set of representative samples. Unfortunately, these approaches can be applied only to specific platforms where large numbers of representative samples have been collected. But there are a large range of platforms for measuring gene expression in use by researchers [13]. Single sample normalization methods are not currently available for many of these platforms.

Even if these normalization methods were available, public measures of gene expression are frequently pre-processed using a range of methods for cleaning, normalization, and analysis, resulting in a range of expression values for the same gene across different platforms [14]. Although these datasets are easily accessible and meta-analyses have been shown to be very powerful [15, 16], platform-specific differences make it difficult to easily and reliably combine cross-platform datasets for such a purpose.

As a concrete example, we focus on the PAM50 signature for breast cancer subtyping [17] which is used to assign patients to one of five breast cancer molecular subtypes: Basal, Luminal A, Luminal B, Her2, Normal. We show that when the number of patients in the test set changes, the predictions for a single patient may change dramatically. We also show that variation in patient populations being predicted on also leads to test set bias.

We then discuss rank-based classifiers, without cross-sample normalization, as an alternative strategy to clinical prediction with genomic data. This approach avoids test set bias by eliminating cross-platform normalization. At the same time, the rank-based approach reduces the influence of platform-specific effects on measurements of gene expression. We show that rank-based subtyping has comparable cross-platform performance with and without normalization.

The simplest rank-based algorithms for prediction are based on pair-wise comparisons of sets of features [18]. We apply the algorithm used to create the PAM50 signature to predict histological grade and compare the approach with a strategy based on predicting with a small number of pairwise gene expression comparisons. We show that both algorithms produce comparable cross-platform performance. We show that the pair-based predictor is both easier to understand and more economical in terms of the number of genes used for prediction. We then develop a 5-pair based replacement for the PAM50 signature and evaluate its performance across 6,297 patients samples from 28 curated breast cancer microarray datasets spanning 15 different platforms curated in inSilicoDb [19, 20]. Our results show that simple rank-based predictors without normalization avoid test set bias and produce easy to interpret, economical, and platform-robust signatures.

## Methods

### Pre-processing and normalization

Pre-processing of the data consisted only of mapping the probe identifiers to NCBI Entrez Gene identifiers so that the probe annotation were consistent across all datasets. Probes were matched to their corresponding gene identifiers using jetset for Affymetrix platforms [21] while for the other platforms, probes with the most variant expression values were chosen for each gene, as previously published [20]. The number of unique genes ranged from ~8,000 to ~20,000 across our compendium of 28 curated datasets. In total, 29,124 genes were represented in at least one platform and 1,531 genes were represented across all technologies (the latter number is restricted by the size of low-density custom-made spotted arrays). Affymetrix datasets were normalized using frozen RMA [22] while, for the other platforms, the gene expression data were analyzed as published; these datasets were originally processed using different normalization approaches by different groups before the dataset was made publicly available. The lack of standardized normalization is representative of data that are available from publicly-available gene expression databases [13].

### Feature creation and classification scheme

Our approach is an extension of the top-scoring pairs and k-top-scoring pairs approaches. The original TSP method suggested one pair of genes that best differentiated classes [18]; k-TSP extended that to the top *k* pairs [23]. We calculated features based on pairwise comparisons of the gene expression levels of genes. Each pair was coded as zero if Gene A *<* Gene B and one if Gene A ≥ Gene B. Models were fit using classification and regression trees [24], resulting in a decision-tree based subtype classifier. Additional information on our model building approach is available in the **Supplementary Information**.

### Feature selection

As we propose in our study to build a robust alternative to the PAM50 subtype classifier published by Parker *et al.* [17], we restricted our analysis to only probes that were present on the platform in question and were part of the original PAM50 subtyping signature. This choice was made to reduce the space of possible pairs, to make the comparison to the PAM50 signature more direct, and to allow the signature we develop to be directly applied to any sample subtyped with a PAM50 assay. Features were selected for the model using a greedy sequential algorithm based on classification and regression trees [25]. For each model fit, we divided the training data into a building and evaluation set. At step one, our algorithm fit a classification and regression tree (CART) model including each of the 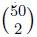 = 1225 possible pairs as a single predictor. The pair with the best error rate on the evaluation set was included in the model. We next considered all models combining the first selected pair and all remaining pairs, and chose the second pair to be the pair that maximally improved accuracy in combination with the first selected pair. This process was continued until five pairs were selected.

### Model trees to account for missing data across platforms

Some genes that are measured on one platform are not measured on others. When predictors are built on a single platform, some of the features that are selected may not be present in the validation sets. To deal with the case when an entire pair is missing in a data set due to one or both of the genes not appearing in a particular platform, we used a model tree approach. We first determine the total set of *N* features estimated from our model selection process. Next, we fit 2*^N^* − 1 models assuming all different possible patterns of missing genes (excluding models with fewer than two features). When attempting to predict on the target dataset, we use the model with the most available pairs. If all pairs are available this results in the default predictor. Further details and analysis appear in **Supplementary Information**.

### Concordance and accuracy measures

We measured the quality of our predictions on the basis of two separate measures: concordance with PAM50 subtypes and accuracy in predicting known pathology of the tumors. Since the PAM50 classifications are themselves predictions, we also examined surrogate measures of the efficacy of our model. ER status and histological grade is known to align with molecular subtypes, although the relationship is not one-to-one [20]. We compared the prognostic value of the PAM50 classification and our rank-based subtype classifier to further assess the biomedical relevance of our new predictive framework.

### Effect of test set size on classification without retraining

When applying predictors on test sets across platforms, it is common to retrain the classifier or to re-normalize the test set to a common scale with the training set. However, in clinical practice, samples will be evaluated one at a time. We estimated the effect of evaluating predictors on sequentially collected test sets through a simulated case scenario where the test set arrived in groups of samples of of varying size.

### Chimeric training set

We established a common gene ID nomenclature across all platforms based on Entrez gene identifiers. Using these IDs we merged samples from multiple platforms and experiments to create chimeric training sets. We then built predictors using pairwise comparisons among the PAM50 genes on these chimeric data sets. Evaluation was performed out-of-sample on additional data both from experiments included in the training set and from completely independent experiments. We describe the value added by this approach in **Supplementary Information**.

### Reproducible research

All curated gene expression and clinical data are available through the InSilicoDb and are programatically available according to the standards of reproducible research [26]. All analyses were performed in the R programming language using Bioconductor [27] packages. The analysis is fully reproducible and can be replicated exactly using the supplementary data, code, and R markdown files [28]. The code to run this analysis is available at http://www.github.com/prpatil/testsetbias.

## Results

### Study population

We collected and curated gene expression microarray data representing 28 independent studies [29]. These datasets spanned 15 different proprietary platform types and a variety of platform versions and included a range of commercial and private manufacturers, spanning Affymetrix, Illumina, and Agilent as well as custom arrays. The data were collected from the Gene Expression Omnibus (GEO) [13], ArrayExpress [30], The University of North Carolina at Chapel Hill database (UNCDB), Stanford Microarray Database (SMD), and Journal and Authors’ websites. Metadata was manually curated as previously described [29]. Experiments ranged from 43 to 1,992 patients, with a median of 131 patients and a total of patients subjects (Table 1).

**Table 1:** Baseline characteristics of curated dataset. Abbreviations: ER - estrogen receptor status; Her2 - human epidermal growth factor receptor 2 status; Node - whether or not cancer has spread to lymph nodes; PGR - progesterone receptor status; RFS - recurrence-free survival time. Age, RFS, Tumor Size are given as means with standard deviations. ^1^due to the ambiguity of grade 2, we chose to build all prediction models for grades 1 and 3 only. ^2^subtypes as predicted by PAM50 with scaling.

| Characteristic | Summary |
| --- | --- |
| n | 6,297 |
| Age (years) | 57.29 (13.42) |
| RFS (years) | 7.22 (4.86) |
| Tumor Size (cm) | 2.52 (1.43) |
| Node |  |
| + | 1,871 |
| − | 2,857 |
| NA | 1,569 |
| Grade<sup>1</sup> |  |
| 1 | 525 |
| 2 | 1,642 |
| 3 | 2,226 |
| NA | 1,904 |
| ER |  |
| + | 3,635 |
| − | 1,556 |
| NA | 1,106 |
| PGR |  |
| + | 766 |
| − | 656 |
| NA | 4,875 |
| Her2 |  |
| + | 496 |
| − | 1,437 |
| NA | 4,364 |
| Subtype<sup>2</sup> |  |
| Basal | 1,254 |
| Her2 | 927 |
| LumA | 2,007 |
| LumB | 1,813 |
| Normal | 296 |

### Normalization makes patient predictions depend on other patients’ data

Consider the PAM50 signature [31]. The class assignment for a new patient is made by calculating a measure of closeness between the new patient and the average patient profile in each class, then choosing the class that was closest to the sample. For example, PAM50 consists of 50 genes and predicts five classes, so each class centroid is a profile of the average expression of each of the 50 genes. The authors used correlation as a measure of closeness for a given sample to each class centroid. This is the step that necessitates suitable rescaling of the test data before predictions are made.

We considered two scenarios which illustrate how PAM50 can produce varying subtype predictions for a single patient if you change the data for other patients used in normalization. We used data from GSE7390 (n = 198), an experiment conducted using the Affymetrix hgu133plus2 microarray. In each experiment, we normalized the gene expression measurements in the test set to fall between 0 and 1.

First we created predictions where we normalized all patients together. Then we calculated predictions for the same patients when normalized in smaller groups (n = 2,10,20,40,80,100,120) and measured the concordance between the predictions for the exact same sample between these two analyses. When normalized in small batches, the predictions for the same patient changed compared to the case where all patients were normalized together (Figures 2).

**Figure 2:**
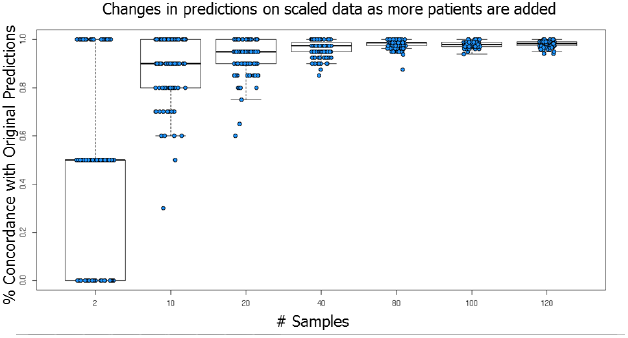
This plot establishes that the predictions for an individual or population can change depending on how many patients are included in the normalization step. We first predicted the PAM50 subtype on an entire set of patients (Affymetrix hgu133plus2; GSE7390; n = 198). We then took 100 random samples of patient subsets ranging from 2-120 patients and predicted their subtypes with data normalization. We compared this newly predicted subtype to each patient’s originally predicted subtype and calculated agreement. Actual data are jittered and overlayed on the boxplot. We find that there is significant variation in percent concordance when a small subset of patients is subtyped in comparison to the entire patient population.

Next we predicted on patient populations that varied in the distribution of ER status. Again we applied the PAM50 predictor to the entire test set. Then we created subsets of the entire test set with differing numbers of ER-negative patients and applied the predictor to each subset. When the percentage of ER-negative patients in the subset matched the percentage in the entire test set, patient subtypes best agreed with the original predictions on the entire test set. However, when the ER status of the other patients in the test set varied, the predictions for the same patient were often different (Figures 3).

**Figure 3:**
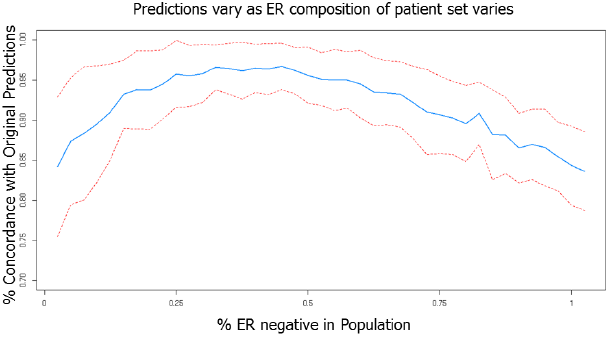
This plot establishes that the predictions for an individual or population can change if the patient population in the test set changes. We first predicted the PAM50 subtype on an entire set of patients (Affymetrix hgu133plus2; GSE7390; n = 198). We then took 100 random samples each of 40 patients and varied the percentage of ER-positive and ER-negative patients in the sample. That is, 0% on the X-axis corresponds to 0% (0/40) ER-negative patients and 100% (40/40) ER-positive patients in the sample. We then predicted subtype on this subset and compared these newly predicted subtypes to the original predictions. The average concordance is plotted with +/- 1SE bands. We note that the original population is 32% ER-negative, which is where we see close to maximal concordance.

### For PAM50 predictions using gene ranks, ignoring normalization is platform-robust

When PAM50 was proposed, the authors chose to calculate similarity based on Spearman correlation [31]. Spearman correlation calculates the correlation between the *ranks* of the two sets of gene expression measurements rather than correlation between the actual values. We hypothesized that this rank-based prediction would be immune to some changes of scale across platforms and other platform-specific artifacts.

To evaluate this hypothesis we used the previously proposed PAM signature building procedure to build a genomic signature for tumor grade again using GSE7390 (n = 198). We built the model on samples measured with Affymetrix and examined the accuracy across other major microarray platforms like Agilent (ISDB10845; n = 337) and Illumina (ISDB10278; n = 1,992).

To predict, we used Spearman correlation in both cases to mimic how the PAM50 signature is used [31]. We predicted new patient samples using our PAM signature for grade both with (red) and without (blue) normalization. We observed that the normalized and un-normalized predictions performed similarly across platforms (Figures 4).

**Figure 4:**
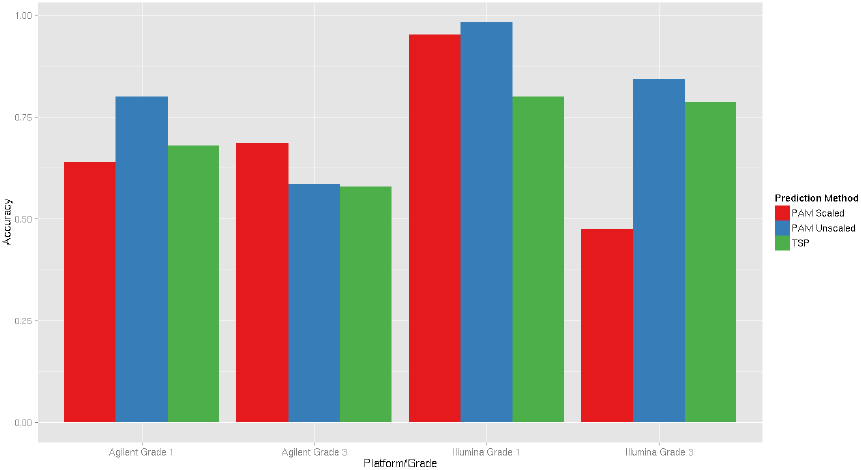
Comparison of accuracy by tumor grade for each of PAM normalized, PAM unnormalized, and the 5-pair TSP model. We see that all three models vary amongst a similar range in their ability to correctly classify tumor grade. We view this as evidence that the purely rank-based procedures (PAM50 unscaled and TSP) perform comparably to a PAM model applied in the traditional manner.

If we do not perform normalization, each patient’s prediction will depend only on their own data. Since the rank based PAM approach performs similarly without normalization, we propose to use unnormalized, rank based approaches for building prediction signatures.

### Simpler rank-based subtyping with pairwise comparisons

PAM50 calculates a prediction by computing the distance from the observed ranks for fifty genes to the centroid ranks for each potential class. It then assigns a patient to the class with the highest correlation. Visually this means that the relative ranks for the fifty genes are plotted against the ranks for each centroid (Figure 5) and the ranking that shows the most correlation is selected. In the example in Figure 5 the patient would be assigned to the Luminal A subtype since their sample was most correlated with the gene centroid for the Luminal A class. But it remains unclear how each gene is contributing to the choice.

**Figure 5:**
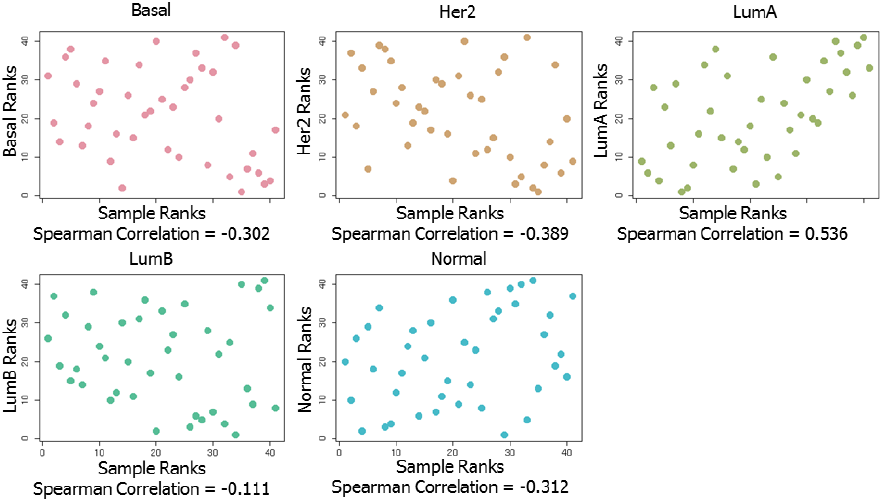
This is a visual description of how the predicted subtype would be chosen for a single patient by the PAM50 predictor. First, the ranks of each of the PAM50 genes in a single patient are compared with the ranks of the PAM50 centroids for each class. For each subtype centroid, the Spearman correlation is calculated. This patient would be classified to Luminal A, as the patient’s gene expression profile is most correlated with the gene centroid for that class.

We applied the top-scoring pair modeling approach to simplify this signature [18]. We searched for pairs of features whose relative ranking would predict which class the patient should be assigned to (as detailed in the **Methods** section) to predict tumor grade. In both this setting and later with intrinsic tumor subtype, we chose to build a chimeric data set consisting of samples from multiple platforms to train the model (n = 1,248 over 8 datasets). The goal was to increase the training sample size and improve the accuracy of our predictor (**Supplementary Information**).

We took the same approach for building a PAM50 substitute predictor as we did when we built a tumor grade predictor above, except we trained our five-pair TSP model on existing PAM50 predictions. We also limited our analysis to the fifty genes that comprise PAM50, primarily so that our predictor could be applicable anywhere where the PAM50 would be used. Our final five-pair TSP decision tree model is shown in Figure 6. In this model it is easier to understand how each gene contributes to the prediction for a patient. At each node, we check whether or not the given gene pair relationship is true in that patient. If it is true, we go right from the node and check the next relationship. If false, we go left.

**Figure 6:**
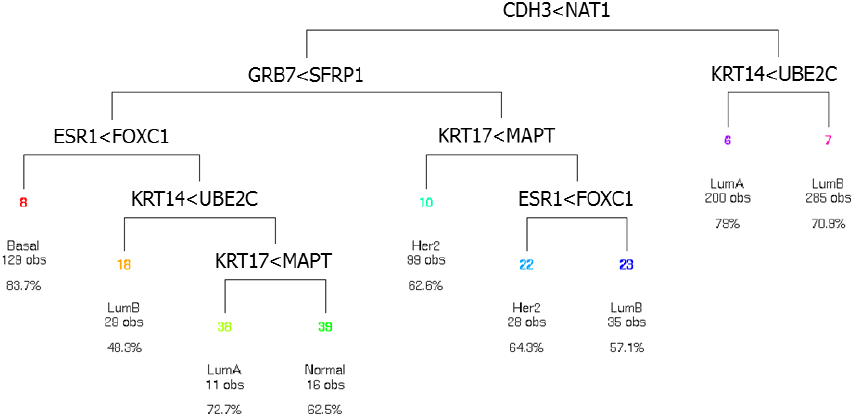
The final decision tree of 5-pair TSP model for intrinsic tumor subtype. With a decision tree, one can examine exactly how a subtype was calculated for a patient based on their gene expression profile. Each node in the tree checks which gene in a pair is expressed higher than the other. The next node to the right or left is chosen based on the result of the previous pairwise relationship, and this is repeated until a subtype is predicted. It is easier to understand how each gene contributes to the eventual prediction.

### Evaluating pair-based signatures

When we used the same procedure to build a pair-based predictor for tumor grade we observed very similar accuracy results for tumor grade as compared with the PAM scaled or unscaled models (Figure 4). After building the TSP model on the chimeric training set as described in the methods section, we applied the TSP predictor to patient samples measured on a range of platforms. We compared our predictions with the original, scaled PAM50 predictions. Here the measure is not accuracy of subtype, but concordance with the original PAM50 predictions. Our predictions show agreement with the original PAM50 predictions amongst the major Affymetrix, Illumina, and Agilent platforms (Figure 7). Older platforms and data set showed more discrepancies.

**Figure 7:**
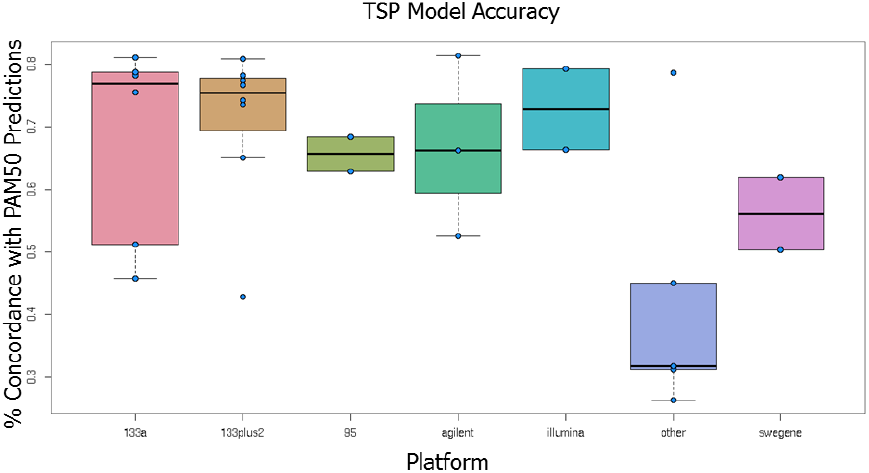
Boxplots of concordance between subtypes predicted by the 5-pair TSP and by the original, scaled PAM50 across platforms. The predictor was built with a chimeric mixture of data from multiple platforms. Each dot represents the accuracy comparing this chimeric TSP to the original, scaled PAM50 classification for one study. The 5-pair TSP agrees fairly well across the three major microarray platforms (Affymetrix, Agilent, Illumina).

Next we compared the ability of the subtypes predicted by PAM50 normalized, PAM50 unnormalized, and our five pair predictor to differentiate survival. We applied all three predictors to all samples (n = 6,297 over 28 platforms). We found that PAM50 predicted subtypes and 5-pair TSP subtypes are informative for survival (**Supplementary Information**). The hazard ratios calculated by Cox proportional hazards models show that all three models are capable of differentiating survival, with similar hazard ratios across all three approaches (Figure 8).

**Figure 8:**
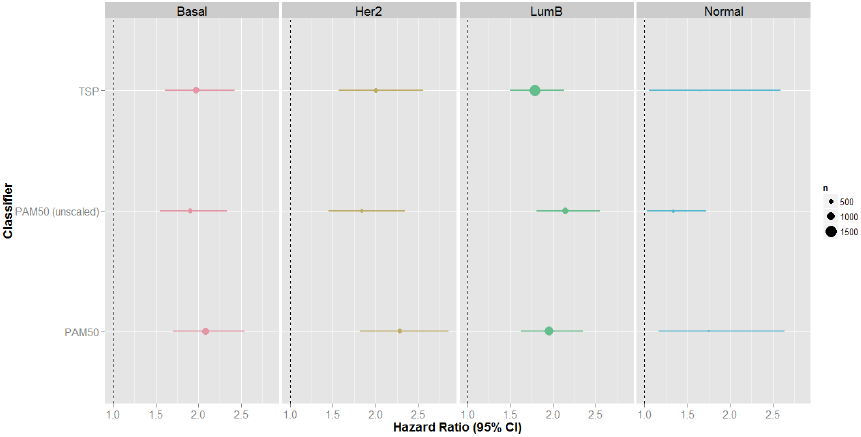
Forest plot of hazard ratios and 95% CIs for each subtype under each classifier. Points are proportional to the sample size in each classified subtype, and Luminal A is used as the reference category for each classifier. Each hazard ratio is significantly different from one, and each subtyper differentiates survival in each subtype comparably.

## Conclusion

Applying PAM50 without scaling is a simple way to avoid the test set bias that can befall any traditional prediction method that depends on data scaling and normalization. We demonstrate that modeling with rank-based features directly maintains this advantage and can additionally produce a simpler and more interpretable model that performs comparably well. Since a key function of PAM50 subtyping is to differentiate survival time amongst patients, the similarity of the hazard ratios in Figure 8 confirms that our TSP-based model produces comparable and useful results.

Other endeavors to reduce the PAM50 signature while maintaining accuracy, such as the three-gene SCM [29], remain completely dependent on data normalization. Here we created a simple 5-pair replacement for PAM50 that can be applied to any sample that has already been assayed with the PAM50 technology. Our predictor has the advantages that (1) it is designed to be applied without normalization and does not suffer from test set bias, (2) it uses fewer genes than the PAM50 signature, and (3) it is easier to interpret the relationship between individual gene expression measurements and the predicted class (Figure Figure 6). Our assay performs similarly in terms of both subtyping concordance and survival differentiation to both the original PAM50 signature both with and without normalization.

## Acknowledgements

This study used data generated by METABRIC; we thank the British Columbia Cancer Agency Branch for sharing these invaluable data with the scientific community.

